# The neural bases of program comprehension: a coordinate-based fMRI meta-analysis

**DOI:** 10.1101/2021.04.15.439937

**Authors:** Yoshiharu Ikutani, Takeshi D. Itoh, Takatomi Kubo

**Affiliations:** Graduate School of Science and Technology, Nara Institute of Science and Technology, 8916-5 Takayama-cho, Ikoma, Nara, 630-0192, Japan

## Abstract

The understanding of brain activity during program comprehension have advanced thanks to noninvasive neuroimaging techniques, such as functional magnetic resonance imaging (fMRI). However, individual neuroimaging studies of program comprehension often provided inconsistent results and made it difficult to identify the neural bases. To identify the essential brain regions, this study performed a small meta-analysis on recent fMRI studies of program comprehension using multilevel kernel density analysis (MKDA). Our analysis identified a set of brain regions consistently activated in various program comprehension tasks. These regions consisted of three clusters, each of which centered at the left inferior frontal gyrus pars triangularis (IFG Tri), posterior part of middle temporal gyrus (pMTG), and right middle frontal gyrus (MFG). Additionally, subsequent analyses revealed relationships among the activation patterns in the previous studies and multiple cognitive functions. These findings suggest that program comprehension mainly recycles the language-related networks and partially employs other domain-general resources in the human brain.

## 1 Introduction

Trillions of lines of source code are running to provide countless services and functionalities essential in our daily life. Software developers expend a vast amount of their working time comprehending existing source code to keep the system maintained and updated (Xia et al., 2017). Thus, many researchers have been investigating the cognitive mechanism of program comprehension as a vital process in software development, in which developers actively acquire knowledge about a software system by reading and exploring program source code (see Storey (2005) and Siegmund (2016) for review papers). This research domain—often referred to as *the neuroscience of programming*—has gained increasing attention from not only software engineering researchers but also cognitive psychologists and neuroscientists (Fedorenko et al., 2019; Rule et al., 2020). In the last decade, noninvasive neuroimaging techniques, such as functional magnetic resonance imaging (fMRI), have advanced our understanding of brain activity during program comprehension. However, individual neuroimaging studies of program comprehension often provided inconsistent results. One of the central contradictions is that multiple studies argued cognitive similarity between program comprehension and natural language comprehension (Siegmund et al., 2014, 2017; Prat et al., 2020; Ikutani et al., 2020), while others emphasized the remarkable difference between them (Floyd et al., 2017; Krueger et al., 2020; Ivanova et al., 2020; Liu et al., 2020).

Previous neuroimaging studies of program comprehension used a variety of experimental tasks with different programming languages. For example, Siegmund et al. (2014, 2017) used Java code snippets and contrasted brain activity during program output estimations against syntax error searches. Ivanova et al. (2020) investigated brain activities measured while subjects read short programs written in Python or ScratchJr and contrasted those activities with content-matched sentence problems written in natural language. Such variability in adopted tasks across individual experiments makes it difficult to evaluate the brain regions consistently activated during program comprehension. Besides, individual neuroimaging studies suffer from limited statistical power and potentially embrace relatively high false-positive rates (Wager et al., 2007; Yarkoni et al., 2010). Therefore, an integrated meta-analysis is necessary to achieve consensus regarding the relationship between brain activity and program comprehension.

Relating program comprehension to diverse human cognitive functions can give a clue to interpret the programming-related brain activity. Since programming language is a relatively new invention in human history, many researchers expect that program comprehension recycles pre-existing cognitive functions in the human brain. Recently, two representative hypotheses are actively studied. One suggests that program comprehension recycles the neural basis of natural language comprehension (Fedorenko et al., 2019). Several fMRI studies support this hypothesis by demonstrating that language-related regions, such as the Broca’s area (BA44/45) and left middle temporal gyrus (MTG), become activated during program comprehension (Siegmund et al., 2014, 2017; Peitek et al., 2021). The other suggests that program comprehension is supported by a frontoparietal network associated with symbol systems (i.e., logic and math) or domain-general executive functions. Two recent studies present evidence supporting this hypothesis by directly comparing activation maps elicited by program comprehension and other cognitive tasks (Ivanova et al., 2020; Liu et al., 2020). A meta-analysis would contribute to evaluating these representative hypotheses and seeking cognitive similarities between program comprehension and other human skills by bridging neuroimaging evidence obtained from program comprehension studies and diverse cognitive/psychological experiments.

Here, we aim to integrate neuroimaging evidence reported in previous program comprehension studies to identify the brain regions consistently activated during program comprehension. To this end, we conduct a coordinate-based meta-analysis of fMRI studies on program comprehension using the multilevel kernel density analysis (MKDA) method. A quantitative meta-analysis is widely used to evaluate the consistency and replicability of activated brain regions across different laboratories, scanners, and task variants. Many peak coordinate-based meta-analyses have been conducted to investigate a variety of cognitive functions and disorders such as working memory (Nee et al., 2013), cognitive control (Derrfuss et al., 2005), and depression (Iwabuchi et al., 2015). This study conducts the first meta-analysis on fMRI studies of program comprehension, to the best of our knowledge. It provides integrated evidence of which regions are essential in a programmer’s brain working for program comprehension.

In addition to the meta-analysis, we seek for relationships between program comprehension and other diverse cognitive functions. To this end, we perform a reverse-inference analysis using a massive database of neuroimaging results. In particular, we employed the NeuroSynth framework (https://neurosynth.org/) developed for large-scale automated meta-analysis of human functional neuroimaging data (Yarkoni et al., 2011). The framework offers a systematic way to identify relevant cognitive functions or states from observed brain activity patterns in a quantitative manner. Inspired by the NeuroSynth and a recent study in cognitive neuroscience (Nakai and Nishimoto, 2020), we investigate voxel-wise similarities between activation patterns of program comprehension tasks and reverse-inference maps associated with broad psychological terms in the NeuroSysnth database. This analysis offers a systematic evaluation procedure to identify a set of cognitive functions potentially related to the ability of program comprehension.

## 2 Materials and Methods

### 2.1 Study selection

To identify fMRI studies on program comprehension, we searched in Google Scholar (https://scholar.google.com/) and PubMed (https://pubmed.ncbi.nlm.nih.gov/) through the Publish or Perish software (https://harzing.com/resources/publish-or-perish) with keywords `[‘program comprehension’ OR ‘code comprehension’] AND [‘fMRI’ OR ‘magnetic resonance imaging’]`. We limited the search scope by publication years from 2011 to 2021 because the research field has started in this decade. We found 276 candidate articles by this search on March 3, 2021. Then, we selected only fMRI studies from these original candidates and excluded the papers reporting experimental results with electroencephalogram (EEG) or near-infrared spectroscopy (NIRS). As a result, we identified only 13 papers describing the results of fMRI experiments in program comprehension. We reviewed the reference lists of all identified papers to avoid missing potentially relevant papers, but we found no addition. To conduct a coordinate-based meta-analysis, we carefully checked the contents of the identified papers and extracted seven papers with at least one table reporting peak voxel coordinates or dataset to replicate the experimental results. As a special case, we included (Siegmund et al., 2017) because the authors provided a table of peak voxel coordinates upon our request.

### 2.2 Multilevel kernel density analysis (MKDA)

We employed the MKDA (Nee et al., 2007; Kober et al., 2008; Wager et al., 2009) for meta-analysis of multiple fMRI studies on program comprehension. The MKDA has been used in many coordinate-based meta-analyses on various psychological domains; e.g., bodily self-awareness (Salvato et al., 2020), moral cognition (Fede and Kiehl, 2019), subjective value (Bartra et al., 2013), and empathy (Fan et al., 2011). Neuroimaging studies typically reported peak voxel coordinates in a particular statistical contrast map (SCM). For instance, Siegmund et al. (2014) derived the peak voxel coordinates from an SCM of blood oxygen level dependent (BOLD) activations in program output estimations against syntax error searches. The MKDA procedure starts with the construction of activation indicator maps from the reported coordinates of each study. The procedure then summarizes the resultant indicator maps with a weight for each study based on its sample size, so that studies with large sample sizes influence more on the meta-analysis. MKDA treats an SCM or a study as the unit of analysis. It produces several advantages over conventional voxel-wise meta-analytical methods (Kober et al., 2008; Wager et al., 2009) which summarise peak coordinates directly (Turkeltaub et al., 2002; Eickhoff et al., 2012). First, the method avoids overestimated statistical power assigned to multiple nearby peaks reported in a study or an SCM. Second, the method can naturally take the sample size and study quality of each SCM into account. Third, the MKDA statistics have a straightforward interpretation: The weighted proportion of SCMs indicates a peak voxel activation within a pre-defined radius of each voxel.

Peak voxel coordinates reported in SCMs were first collected from the selected fMRI studies of program comprehension (see Figure 1a for a procedural overview). Technically, the voxel coordinates were described in one of two stereotaxic spaces: the Talairach space or Montreal Neurological Institute (MNI) space. We used the MNI space as the reference and converted all maps with Talairach coordinates to MNI space prior to the analysis. The peaks obtained from each SCM were convolved with a spherical kernel with a 15 mm radius and thresholded at a maximum value of 1, so that multiple nearby peaks were not counted as multiple activations. This kernel convolution yielded an indicator map for each SCM, where a voxel value of 1 indicates the existence of a peak within a preset radius while 0 indicates the absence of peaks. A weighted average of the resulting indicator maps was then calculated to provide an interpretable metric; the assigned weight of each SCM was defined as the square root of the sample size in each study. Finally, significant regions were determined using a statistical threshold obtained through 10,000 iterations of a Monte-Carlo simulation. The null hypothesis was that the convolved peaks in the indicator maps were not spatially consistent, or distributed randomly throughout the brain. We report the thresholded results as an MKDA statistic map with a voxel-level threshold of p ¡ 0.001 (uncorrected) and a cluster threshold of p ¡ 0.05 (family-wise error [FWE]-corrected). Note that our meta-analysis included the peak voxel coordinates reported in Ikutani et al. (2020), derived from decoding analysis but not a BOLD contrast, to cover as many available neuroimaging results as possible in the current literature.

**Figure 1:**
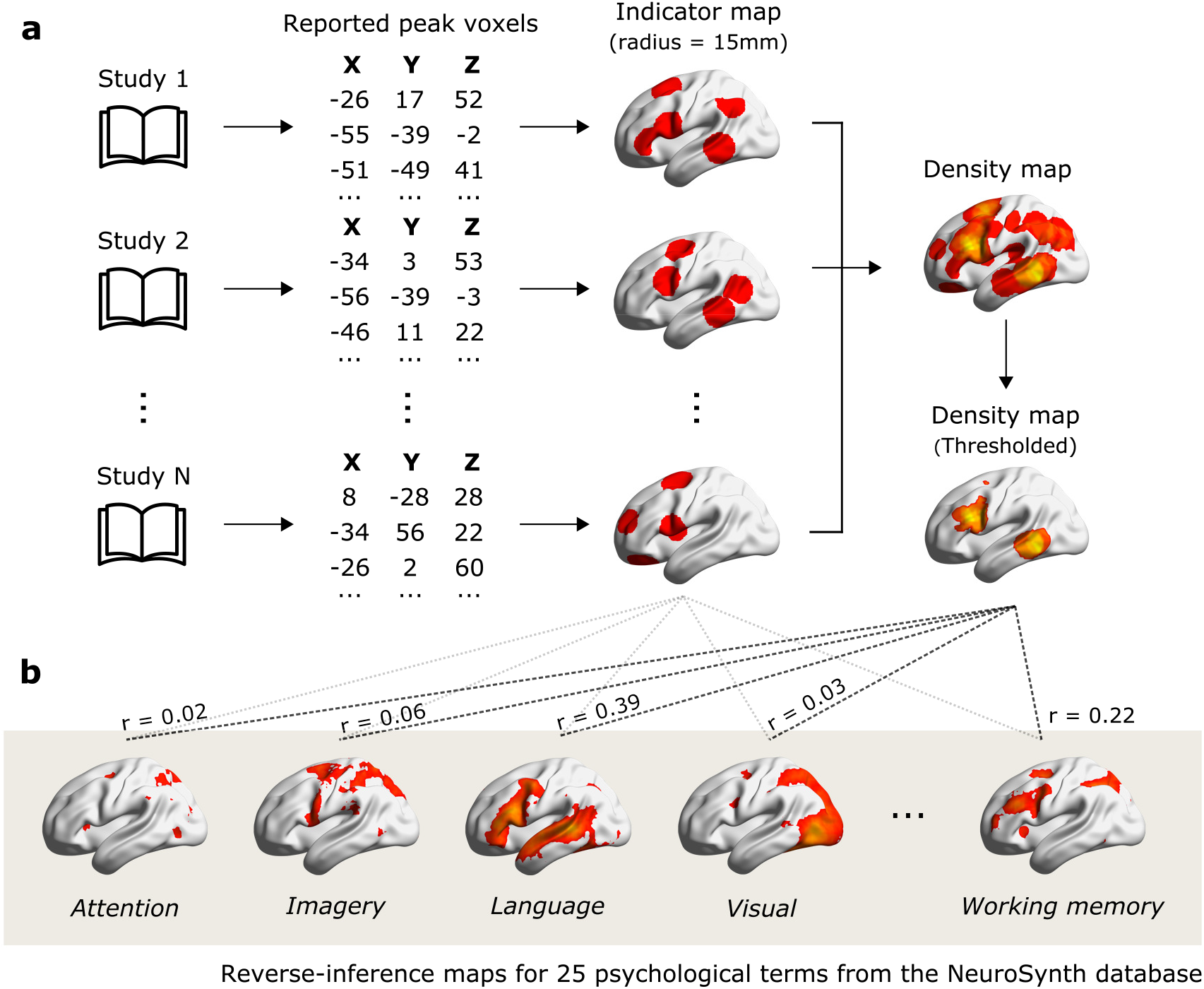
Overview of the meta-analysis methods. (**a**) Schematic chart of multilevel kernel density analysis (MKDA). Indicator maps were obtained by convolving peak voxel coordinates with a spherical kernel with 15 mm radius. Significant threshold for the density map was determined by a Monte-Carlo simulation. (**b**) Metadata-based evaluation of MKDA results. To quantify voxel-wise cognitive similarity, Pearson’s correlation coefficients were calculated between the MKDA resultant maps and the reverse-inference maps taken from the NeuroSynth database.

### 2.3 Metadata-based evaluation of MKDA results

To relate program comprehension to diverse human cognitive functions, we performed a metadata-based evaluation of our MKDA results. We selected 25 psychological terms used in Yarkoni et al. (2011) as a reference set of cognitive functions and the reverse-inference maps (also known as association test maps) for each term taken from the NeuroSynth database. Each voxel value in a reverse-inference map indicates the likelihood of the target term occurrence in a study where an activation is observed at the voxel. As a quantitative measure of cognitive similarity, we calculated Pearson’s correlation coefficients of our MKDA resultant maps with each reverse-inference map (see Figure 1b for an overview). We ignored negative values in the reverse-inference maps in this calculation by assigning zero. Additionally, we conducted a hierarchical clustering analysis (HCA) to examine relationships across individual fMRI studies on program comprehension. We used a set of correlation coefficients to each reverse-inference map as a feature vector of each MKDA resultant map. We then constructed a dendrogram of ten maps (one thresholded density map and nine indicator maps) using the feature dissimilarity (1 - correlation coefficient) as a distance metric and the minimum distance as a linkage criterion. Our metadata-based evaluation procedure offers a systematic way to interpret the activation map drawn from multiple fMRI studies of program comprehension. A systematic evaluation is important to achieve consensus across multiple neuroimaging studies supporting different conclusions. Besides, this evaluation procedure moderates interpretation biases caused by motivations and expected outcomes underlying the authors.

## 3 Results

### 3.1 Description of the selected studies

Seven studies with nine contrasts and 61 peak coordinates were included in our meta-analysis (see Table 1 for the list of all studies included). Among these nine contrasts, six contrasts investigated program output estimation against control tasks, and two examined BOLD activations during source code inspections. Peak voxel coordinates in Ikutani et al. (2020) showed significant correlations between decoding accuracy of source code categories and behavioral performance of program categorization tasks. From the the pro-gramming language perspective, four studies used Java, two used Python, and one study each used C and ScratchJr in the experiments. The average and standard deviation of the sample size were 19.50 ± 5.58. The square root of the sample size used as the weights of each SCM in MKDA ranged from 3.32 to 5.48. Peak voxels in five contrasts represented by the Talairach space were converted to MNI coordinates in advance to the subsequent meta-analysis.

**Table 1:** fMRI studies on program comprehension included in our meta-analysis.

| Study | Language | # subjects | Contrast | # peaks | Space |
| --- | --- | --- | --- | --- | --- |
| Siegmund,<br>2014 | Java | 17 | Program output estimation<br>> Control (Syntax error search) | 5 | TAL |
| Siegmund,<br>2017 | Java | 11 | Program output estimation with semantic cues<br>> Program output estimation without semantic cues | 5 | TAL |
| Castelhano,<br>2019 | C | 20 | Inspection of source code<br>> Inspection of pseudo-code text | 8 | TAL |
| Castelhano,<br>2019 | C | 20 | Inspection of source code with bugs<br>> Inspection of source code without bugs | 9 | TAL |
| Liu,<br>2020 | Python | 17 | Program output estimation<br>> Inspection of fake code | 7 | MNI |
| Ivanova,<br>2020 | Python | 24 | Program output estimation<br>> Output estimation from sentence | 9 | MNI |
| Ivanova,<br>2020 | ScratchJr | 19 | Program output estimation<br>> Output estimation from sentence | 7 | MNI |
| Peitek,<br>2021 | Java | 18 | Program output estimation<br>> Control (Locating brackets) | 4 | TAL |
| Ikutani,<br>2020 | Java | 30 | Significant correlation between decoding accuracy<br>of program categories and behavioral performance | 7 | MNI |

### 3.2 Brain regions consistently activated in program comprehension studies

We first investigated which brain regions were consistently activated across program comprehension studies included in our meta-analysis. The MKDA identified three primary clusters at a whole-brain statistical threshold (voxel-level p ¡ 0.001 [uncorrected] and cluster-level p ¡ 0.05 [FWE-corrected]). See Figure 2a for the spatial distribution of reported peaks and Figure 2b for the MKDA result. Table 2 describes the voxel locations of the peak weighted densities in each cluster. The first cluster was centered at the left inferior frontal gyrus pars triangularis (IFG Tri), covering the large part of Broca’s area. The second cluster was centered at the left posterior middle temporal gyrus (pMTG). The third was on the right middle frontal gyrus (MFG) extending to the adjacent right superior and inferior frontal gyri. In contrast to several previous studies (Siegmund et al., 2014; Liu et al., 2020; Ivanova et al., 2020; Peitek et al., 2021), left parietal regions such as angular gyrus and inferior parietal lobule were not observed in our MKDA result at the significant threshold.

**Table 2:**
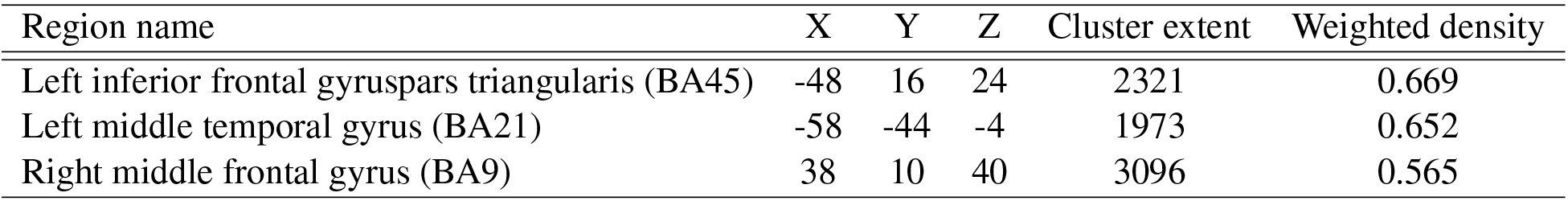
Significant clusters identified by MKDA.

| Region name | X | Y | Z | Cluster extent | Weighted density |
| --- | --- | --- | --- | --- | --- |
| Left inferior frontal gyrus pars triangularis (BA45) | -48 | 16 | 24 | 2321 | 0.669 |
| Left middle temporal gyrus (BA21) | -58 | -44 | -4 | 1973 | 0.652 |
| Right middle frontal gyrus (BA9) | 38 | 10 | 40 | 3096 | 0.565 |

**Figure 2:**
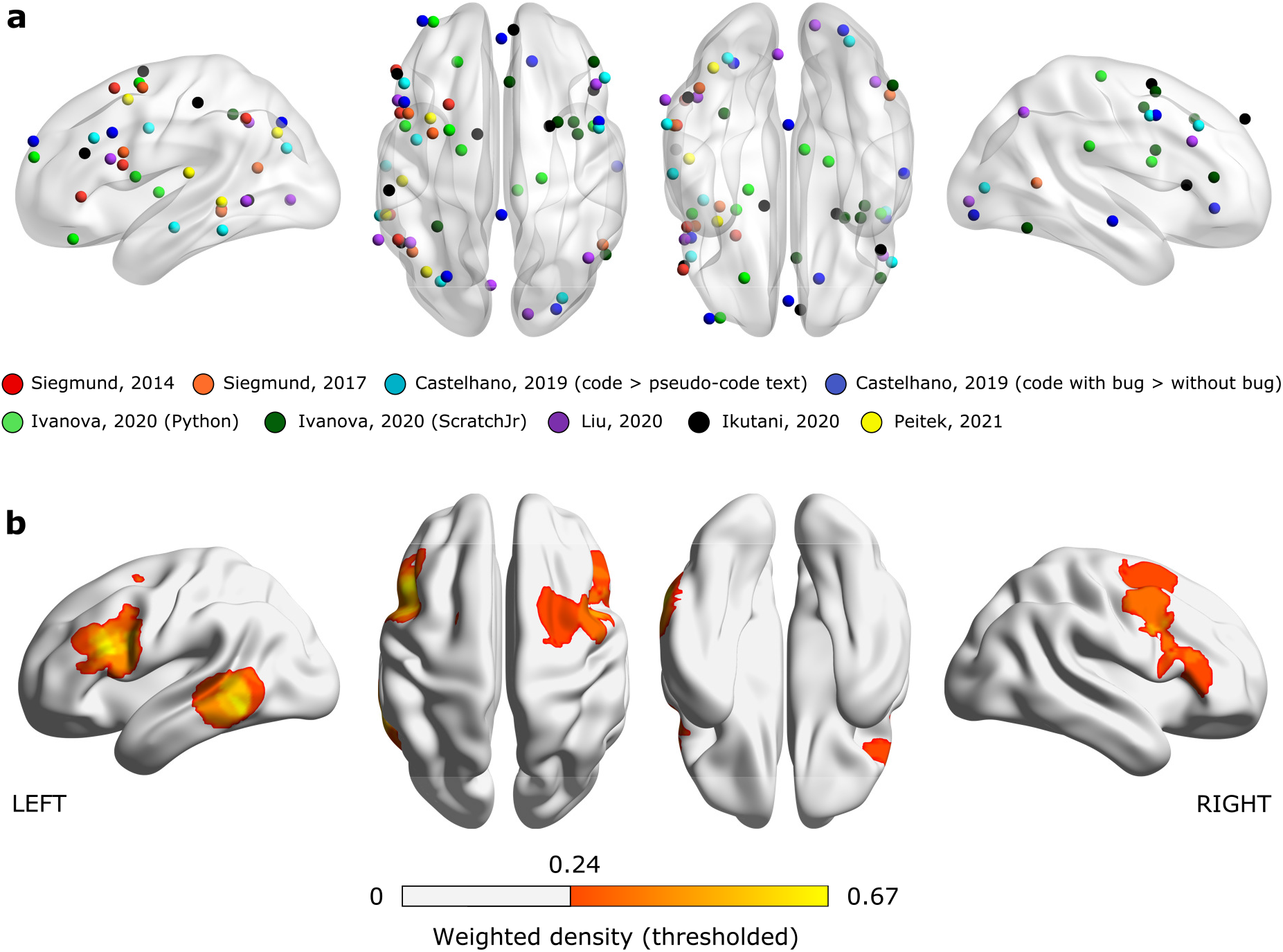
Result of the multi-level kernel density analysis. (**a**) Peak voxel coordinates from the nine contrasts included in the meta-analysis. All peaks were plotted in a single ICBM 152 template brain using the BrainNet Viewer (Xia et al., 2013). (**b**) Brain regions showing consistent activation across all program comprehension studies at a whole-brain statistical thresholds (voxel-level p < 0.001 [uncorrected] and cluster-level p < 0.05 [FWE-corrected]).

### 3.3 Relating program comprehension to diverse human cognitive functions

Our metadata-based evaluation procedure revealed the relationship of program comprehension with diverse human cognitive functions (Figure 3). The resultant density map of our meta-analysis showed significant correlations to the multiple reverse-inference maps. The top-5 largest correlations were observed with the reverse-inference maps generated for *Language*, *Phonological*, *Semantic*, *Working memory*, and *Verbal*. The indicator maps taken from Siegmund et al. (2014, 2017), and Liu et al. (2020) showed similar patterns to the meta-analysis result. The figure also indicates that two indicator maps taken from different contrasts in Castelhano et al. (2019) were located apart from each other in the HCA dendrogram. Specifically, the contrast of program comprehension, that is, [Inspection of source code > Inspection of pseudo-code text] was located closer to the meta-analysis result than the bug searching contrast, [Inspection of source code with bugs > Inspection of source code without bugs]. The indicator maps taken from Ikutani et al. (2020) and Peitek et al. (2021) had slightly different patterns but still showed *Language* and *Semantic* within the top-3 largest correlations. Two indicator maps taken from contrasts of [Program output estimation > Output estimation from sentence] in Ivanova et al. (2020) were grouped by the unique cluster in our HCA dendrogram, having very small correlations to the language-related terms. The indicator map of ScrachJr in Ivanova et al. (2020) showed a different pattern than others. The top-5 largest correlations were observed with *Working memory*, *Conflict*, *Interference*, *Attention*, and *Executive*. Further, the indicator map of Python in Ivanova et al. (2020) showed relatively small correlations to all psychological terms.

**Figure 3:**
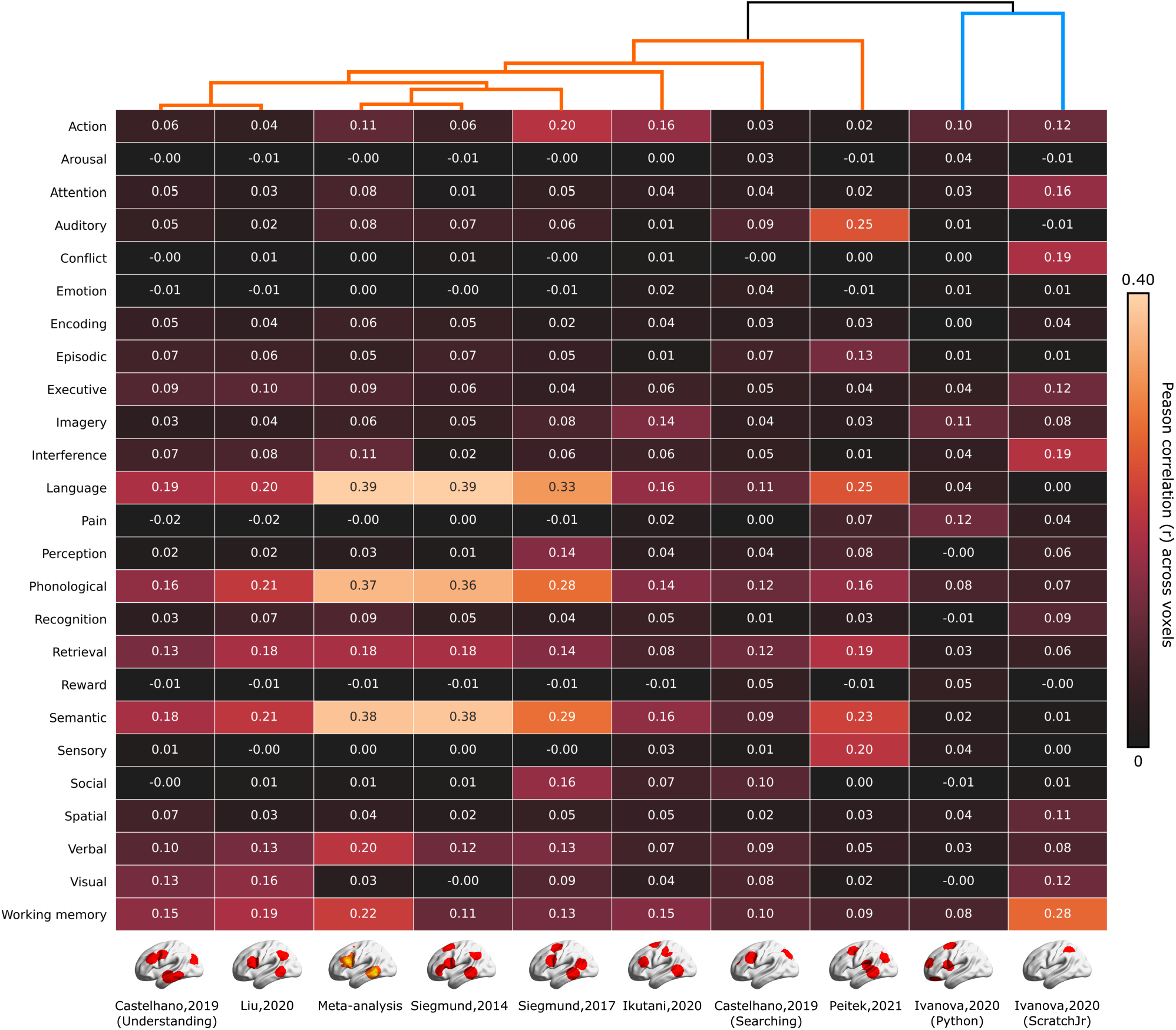
Metadata-based evaluation of the MKDA results. Each cell represents Peason’s correlation coefficients of each indicator map or thresholded density map with the reverse-inference maps for 25 psychological terms used in (Yarkoni et al., 2011). A dendrogram shown in top of the figure indicates the hierarchical organization of individual fMRI studies on program comprehension evaluated using a hierarchical clustering analysis (HCA).

## 4 Discussion

Our quantitative meta-analysis of fMRI studies on program comprehension identified a set of brain regions consistently activated in various program comprehension tasks and stimuli. The region set consisted of three clusters centered at the left IFG Tri, left pMTG, and right MFG, respectively. The subsequent metadata-based evaluation revealed the relationship of our meta-analysis result with the reverse-inference maps generated for language-related terms, such as *Language*, *Phonological*, and *Semantic*. All indicator maps except those taken from Ivanova et al. (2020) showed similar patterns to the meta-analysis result, signaling the functional similarity between program comprehension and natural language comprehension. This study provides the first meta-analysis on fMRI studies of program comprehension, to the best of our knowledge. It evaluates the consistency and replicability of published neuroimaging results associated with various program comprehension activities.

The meta-analytic evidence supports the view that program comprehension recycles the neural basis of natural language comprehension. We demonstrated that the left IFG Tri and pMTG were consistently activated in various program comprehension tasks. The left IFG Tri and the left pMTG are major components of the language network in the human brain and functionally associated with semantic selecting/retrieving tasks (Demonet et al., 1992; Thompson-Schill et al., 1997; Simmons et al., 2005; Price, 2012). In addition, several studies indicated that these two regions are sensitive to cognitive demands for directing semantic knowledge retrieval in a goal-oriented way (Rodd et al., 2005; Kuhl et al., 2007; Whitney et al., 2010). The activations of the left IFG Tri and pMTG were repeatedly found in BOLD contrasts of “real” program output estimations against “fake” code inspections. These contrasts were typically designed to capture how a programmer’s brain works for understanding the contents of source code by subtracting the brain activities elicited from seeing meaningless pseudo-code as a visual stimulus. In other words, these contrasts may emphasize the programmer’s reading behavior to extract the meaning of source code while canceling the shared activations related to sensory processing. Based on our meta-analytic results, programmers’ reading behavior on source code recruits the language network in the brain, suggesting the ability of program comprehension might be realized by recycling the neural basis of natural language comprehension.

Multiple psychological and neuroimaging studies have emphasized the cognitive similarity between program comprehension and natural language comprehension. Siegmund et al. (2014, 2017) contrasted BOLD signals in program output estimations against non-semantic control tasks and reported the activations of language-related regions, that is, the left IFG Tri and pMTG, in their first and subsequent replication studies. A recent decoding-based study found that decoding accuracy of source code categories in the left IFG Tri and pMTG were significantly correlated with behavioral performances in expert programmers (Ikutani et al., 2020). These findings suggest that a programmer’s brain recruits a similar language-related network for both natural language processing and program comprehension. Further, several researchers hypothesized that learning a modern programming language resembles learning a second natural language in adulthood (Fedorenko et al., 2019). A recent study demonstrated that behavioral and EEG-based language-learning aptitude measures explain significant variance in the learning rates of subjects who participated in Python training sessions (Prat et al., 2020). Our meta-analytic evidence may support these perspectives by offering an integrated view across multiple fMRI studies on program comprehension.

In contrast, several previous studies underline the difference between program comprehension and natural language comprehension. Floyd et al. (2017) argued the discriminability of brain activities between program and natural language comprehension (more precisely, code and prose reviews) using decoding-based analysis. Liu et al. (2020) scanned 17 expert programmers and measured brain activities while subjects performed program comprehension as well as localizer tasks for memory control, formal logic, symbolic math, executive control, and language. The study described the functional similarity of program comprehension to formal logical inference by examining the overlap between the activation maps revealed by the code contrast and each of the localizer contrasts. Ivanova et al. (2020) investigated two candidate brain systems, the multiple demand (MD) system and the language system, that potentially supports the ability of program comprehension. Their results indicated that the MD system showed strong responses to source code written in Python and ScratchJr, whereas the language system responded strongly to sentence problems but weakly to source code problems. These controversial results could be explained by viewing program comprehension as a set of micro cognitive processes. We consider that program comprehension requires multiple cognitive processes involving a) read characters and texts, b) associate described information with internal knowledge, and c) infer or reconstruct plausible mental models of how the program works. Our meta-analytic evidence may support that program comprehension recruits the language-related network only for realizing the initial reading process. Non-language brain networks might be recruited for the later cognitive processes potentially associated with knowledge retrieval and/or mental model reconstruction.

Another important finding in our meta-analysis is that the activations in the right MFG are consistently observed across multiple fMRI studies on program comprehension. Specifically, the right MFG activations were reported in six contrasts taken from Castelhano et al. (2019), Ivanova et al. (2020), Liu et al. (2020), and Ikutani et al. (2020). In literature, the right MFG is frequently associated with spatial working memory (McCarthy et al., 1994; Leung et al., 2002) and considered to be a component of the MD system network, which is typically recruited for math, logic, and problem-solving tasks (Fedorenko et al., 2013; Koyama et al., 2017). Our metadata-based evaluation procedure also showed that the MKDA result of our meta-analysis was strongly correlated with the reverse-inference map generated for *Working memory*. This evidence collectively implies that the various program comprehension tasks included in our meta-analysis may recruit the cognitive resources of spatial working memory and/or MD systems. This finding potentially suggests that program comprehension is not a simple derivation of natural language processing, but could be realized by complex orchestrations of the language network and other cognitive resources.

Our meta-analytic framework provides a promising way to achieve the consensus regarding the relationship between brain activity and program comprehension. Evaluating programmer’s brain activities during program comprehension is difficult because many factors (e.g., choice of programming language, task design, or contrast) affect the study outcomes. For instance, Ivanova et al. (2020) contrasted BOLD activations of program output estimations against content-matched sentence problems, while other studies contrasted it against inspections of meaningless pseudo-code. Consequently, their experimental setting subtracted activations of natural language comprehension from the activations during program output estimations. In fact, our meta-analytic results greatly reflected such differences as the indicator maps taken from Ivanova et al. (2020) were grouped into a unique cluster (Figure 3). This example throws out the caveat that neuroimaging evidence obtained in a single study could be biased by its experimental setting and potentially not enough to be generalized for program comprehension using different languages and scopes. Similar to the recent studies in cognitive neuroscience (Huth et al., 2016; Nakai and Nishimoto, 2020), future work with diverse task settings involving different programming languages and behaviors should be expected in the domain of the neuroscience of programming.

### 4.1 Limitations of the study

This meta-analysis included only seven studies with nine contrasts and 61 peak coordinates, indicating its relatively smaller sample size compared to previous MKDA-based meta-analyses. For example, Fan et al. (2011) conducted a meta-analysis of emotion upon 40 studies with 50 contrasts and Salvato et al. (2020) recruited 40 studies with 73 contrasts in a meta-analysis of bodily self-awareness. The shortage of included studies is not surprising because the neuroscience of programming is still in its infancy, and there are only a handful of published fMRI studies in the domain. Specifically, our study selection procedure has identified 13 papers, but only seven papers had with tables describing peak voxel coordinates or datasets to replicate the experimental results. Therefore, the meta-analytic results obtained via this study might be biased due to the small sample size, and the conclusion could be changed if we will get more neuroimaging studies on program comprehension. Making mention of the neural basis of program comprehension must be with caution from the currently available evidence. We encourage authors to open their peak voxel tables and experimental data for future replication and meta-analyses.

Our metadata-based evaluation procedure only considered Pearson’s correlation coefficients across voxels between our MKDA resultant maps and the reverse-inference maps taken from the NeuroSynth database. Furthermore, the procedure systematically ignored the strength and individual differences of BOLD signals reported in each peak voxel because the MKDA only evaluated the locations where peak voxels were observed. Due to these limitations, the results obtained in our meta-analysis might overestimate spatial consistency than other information. Comparing brain activity patterns during diverse task settings within an individual subject would be a more rigorous scheme to investigate the relationship of program comprehension to diverse human cognitive functions. Additionally, differences in brain activity patterns between different programming languages or programming expertise (i.e., experts against novice programmers) should be examined in future work.

## Acknowledgements

We thank J. Siegmund, N. Peitek for sharing their data of peak voxel coordinates in Siegmund et al. (2017) and A. A. Ivanova for helping the replication of two contrast maps in Ivanova et al. (2020). This work was supported by JSPS KAKENHI Grant Numbers JP16H06569, JP18J22957, and JP19J20669.

## References

Oscar Bartra, Joseph T McGuire, and Joseph W Kable. The valuation system: a coordinate-based meta-analysis of bold fmri experiments examining neural correlates of subjective value. Neuroimage, 76: 412–427, 2013.

Joao Castelhano, Isabel C Duarte, Carlos Ferreira, Joao Duraes, Henrique Madeira, and Miguel Castelo-Branco. The role of the insula in intuitive expert bug detection in computer code: an fMRI study. Brain Imaging and Behavior, 13(3):623–637, 2019.

Jean-François Demonet, Francois Chollet, Stuart Ramsay, Dominique Cardebat, Jean-Luc Nespoulous, Richard Wise, André Rascol, and Richard Frackowiak. The anatomy of phonological and semantic processing in normal subjects. Brain, 115(6):1753–1768, 1992.

Jan Derrfuss, Marcel Brass, Jane Neumann, and D Yves von Cramon. Involvement of the inferior frontal junction in cognitive control: Meta-analyses of switching and stroop studies. Human brain mapping, 25 (1):22–34 2005.

Simon B Eickhoff, Danilo Bzdok, Angela R Laird, Florian Kurth, and Peter T Fox. Activation likelihood estimation meta-analysis revisited. Neuroimage, 59(3):2349–2361, 2012.

Yan Fan, Niall W Duncan, Moritz de Greck, and Georg Northoff. Is there a core neural network in empathy? an fmri based quantitative meta-analysis. Neuroscience & Biobehavioral Reviews, 35(3):903–911, 2011.

Samantha J Fede and Kent A Kiehl. Meta-analysis of the moral brain: patterns of neural engagement assessed using multilevel kernel density analysis. Brain imaging and behavior, pages 1–14 2019.

Evelina Fedorenko, John Duncan, and Nancy Kanwisher. Broad domain generality in focal regions of frontal and parietal cortex. Proceedings of the National Academy of Sciences, 110(41):16616–16621, 2013.

Evelina Fedorenko, Anna Ivanova, Riva Dhamala, and Marina Umaschi Bers. The language of programming: a cognitive perspective. Trends in cognitive sciences, 23(7):525–528, 2019.

Benjamin Floyd, Tyler Santander, and Westley Weimer. Decoding the representation of code in the brain: An fMRI study of code review and expertise. In Proceedings of the IEEE/ACM 39th International Conference on Software Engineering, pages 175–186. IEEE, 2017.

Alexander G Huth, Wendy A de Heer, Thomas L Griffiths, Frédéric E Theunissen, and Jack L Gallant. Natural speech reveals the semantic maps that tile human cerebral cortex. Nature, 532(7600):453–458, 2016.

Yoshiharu Ikutani, Takatomi Kubo, Satoshi Nishida, Hideaki Hata, Kenichi Matsumoto, Kazushi Ikeda, and Shinji Nishimoto. Expert programmers have fine-tuned cortical representations of source code. Eneuro, 8(1), 2020.

Anna A Ivanova, Shashank Srikant, Yotaro Sueoka, Hope H Kean, Riva Dhamala, Una-May O’Reilly, Marina U Bers, and Evelina Fedorenko. Comprehension of computer code relies primarily on domain-general executive brain regions. eLife, 9:e58906, 2020. doi: 10.7554/eLife.58906.

Sarina J Iwabuchi, Rajeev Krishnadas, Chunbo Li, Dorothee P Auer, Joaquim Radua, and Lena Palaniyappan. Localized connectivity in depression: a meta-analysis of resting state functional imaging studies. Neuroscience & Biobehavioral Reviews, 51:77–86 2015.

Hedy Kober, Lisa Feldman Barrett, Josh Joseph, Eliza Bliss-Moreau, Kristen Lindquist, and Tor D Wager. Functional grouping and cortical–subcortical interactions in emotion: a meta-analysis of neuroimaging studies. Neuroimage, 42(2):998–1031, 2008.

Maki S Koyama, David O’Connor, Zarrar Shehzad, and Michael P Milham. Differential contributions of the middle frontal gyrus functional connectivity to literacy and numeracy. Scientific reports, 7(1):1–13, 2017.

Ryan Krueger, Yu Huang, Xinyu Liu, Tyler Santander, Westley Weimer, and Kevin Leach. Neurological divide: an fmri study of prose and code writing. In 2020 IEEE/ACM 42nd International Conference on Software Engineering (ICSE), pages 678–690. IEEE, 2020.

Brice A Kuhl, Nicole M Dudukovic, Itamar Kahn, and Anthony D Wagner. Decreased demands on cognitive control reveal the neural processing benefits of forgetting. Nature Neuroscience, 10(7):908, 2007.

H-C Leung, John C Gore, and Patricia S Goldman-Rakic. Sustained mnemonic response in the human middle frontal gyrus during on-line storage of spatial memoranda. Journal of cognitive neuroscience, 14 (4):659–671 2002.

Yun-Fei Liu, Judy Kim, Colin Wilson, and Marina Bedny. Computer code comprehension shares neural resources with formal logical inference in the fronto-parietal network. eLife, 9:e59340, 2020. doi: 10.7554/eLife.59340.

Gregory McCarthy, Andrew M Blamire, Aina Puce, Anna C Nobre, Gilles Bloch, Fahmeed Hyder, Patricia Goldman-Rakic, and Robert G Shulman. Functional magnetic resonance imaging of human prefrontal cortex activation during a spatial working memory task. Proceedings of the National Academy of Sciences, 91(18):8690–8694, 1994.

Tomoya Nakai and Shinji Nishimoto. Quantitative models reveal the organization of diverse cognitive functions in the brain. Nature communications, 11(1):1–12, 2020.

Derek Evan Nee, Tor D Wager, and John Jonides. Interference resolution: insights from a meta-analysis of neuroimaging tasks. Cognitive, Affective, & Behavioral Neuroscience, 7(1):1–17, 2007.

Derek Evan Nee, Joshua W Brown, Mary K Askren, Marc G Berman, Emre Demiralp, Adam Krawitz, and John Jonides. A meta-analysis of executive components of working memory. Cerebral cortex, 23(2): 264–282 2013.

Norman Peitek, Sven Apel, Chris Parnin, André Brechmann, and Janet Siegmund. Program comprehension and code complexity metrics: An fmri study. In Proceedings of the IEEE/ACM 43rd International Conference on Software Engineering. IEEE, 2021.

Chantel S Prat, Tara M Madhyastha, Malayka J Mottarella, and Chu-Hsuan Kuo. Relating natural language aptitude to individual differences in learning programming languages. Scientific reports, 10(1):1–10, 2020.

Cathy J Price. A review and synthesis of the first 20 years of PET and fMRI studies of heard speech, spoken language and reading. NeuroImage, 62(2):816–847, 2012.

Jennifer M Rodd, Matthew H Davis, and Ingrid S Johnsrude. The neural mechanisms of speech comprehension: fMRI studies of semantic ambiguity. Cerebral Cortex, 15(8):1261–1269, 2005.

Joshua S Rule, Joshua B Tenenbaum, and Steven T Piantadosi. The child as hacker. Trends in Cognitive Sciences, 24(11):900–915, 2020.

Gerardo Salvato, Fabian Richter, Lucas Sedeño, Gabriella Bottini, and Eraldo Paulesu. Building the bodily self-awareness: Evidence for the convergence between interoceptive and exteroceptive information in a multilevel kernel density analysis study. Human brain mapping, 41(2):401–418, 2020.

Janet Siegmund. Program comprehension: Past, present, and future. In Proceedings of the 23rd International Conference on Software Analysis, Evolution, and Reengineering, pages 13–20 2016. doi: 10.1109/SANER.2016.35.

Janet Siegmund, Christian Kästner, Sven Apel, Chris Parnin, Anja Bethmann, Thomas Leich, Gunter Saake, and André Brechmann. Understanding understanding source code with functional magnetic resonance imaging. Proceedings of the IEEE/ACM 36th International Conference on Software Engineering, pages 378–389 2014.

Janet Siegmund, Norman Peitek, Chris Parnin, Sven Apel, Johannes Hofmeister, Christian Kästner, Andrew Begel, Anja Bethmann, and André Brechmann. Measuring neural efficiency of program comprehension. Proceedings of the 11th Joint Meeting on Foundations of Software Engineering, pages 140–150 2017.

Alan Simmons, Daniel Miller, Justin S Feinstein, Terry E Goldberg, and Martin P Paulus. Left inferior prefrontal cortex activation during a semantic decision-making task predicts the degree of semantic organization. NeuroImage, 28:30–38 2005.

Margaret-Anne Storey. Theories, methods and tools in program comprehension: past, present and future. In Proceedings of the 13th International Workshop on Program Comprehension, pages 181–191 2005.

Sharon L Thompson-Schill, Mark D’Esposito, Geoffrey K Aguirre, and Martha J Farah. Role of left inferior prefrontal cortex in retrieval of semantic knowledge: a reevaluation. Proceedings of the National Academy of Sciences, 94(26):14792–14797, 1997.

Peter E Turkeltaub, Guinevere F Eden, Karen M Jones, and Thomas A Zeffiro. Meta-analysis of the functional neuroanatomy of single-word reading: method and validation. Neuroimage, 16(3):765–780, 2002.

Tor D Wager, Martin Lindquist, and Lauren Kaplan. Meta-analysis of functional neuroimaging data: current and future directions. Social cognitive and affective neuroscience, 2(2):150–158, 2007.

Tor D Wager, Martin A Lindquist, Thomas E Nichols, Hedy Kober, and Jared X Van Snellenberg. Evaluating the consistency and specificity of neuroimaging data using meta-analysis. Neuroimage, 45(1):S210–S221, 2009.

Carin Whitney, Elizabeth Jefferies, and Tilo Kircher. Heterogeneity of the left temporal lobe in semantic representation and control: priming multiple versus single meanings of ambiguous words. Cerebral Cortex, 21(4):831–844, 2010.

Mingrui Xia, Jinhui Wang, and Yong He. BrainNet Viewer: a network visualization tool for human brain connectomics. PLoS One, 8(7):e68910, 2013.

X. Xia, L. Bao, D. Lo, Z. Xing, A. E. Hassan, and S. Li. Measuring program comprehension: A large-scale field study with professionals. IEEE Transactions on Software Engineering, 44(10):951–976, 2017.

Tal Yarkoni, Russell A Poldrack, David C Van Essen, and Tor D Wager. Cognitive neuroscience 2.0: building a cumulative science of human brain function. Trends in cognitive sciences, 14(11):489–496, 2010.

Tal Yarkoni, Russell A Poldrack, Thomas E Nichols, David C Van Essen, and Tor D Wager. Large-scale automated synthesis of human functional neuroimaging data. Nature methods, 8(8):665–670, 2011.

